# Identifying virulence determinants of multidrug-resistant *Klebsiella pneumoniae* in *Galleria mellonella*

**DOI:** 10.1101/2020.10.30.362657

**Authors:** Sebastian Bruchmann, Theresa Feltwell, Julian Parkhill, Francesca L. Short

## Abstract

Infections caused by *Klebsiella pneumoniae* are a major public health threat. Extensively drug-resistant and even pan-resistant strains have been reported. Understanding *K. pneumoniae* pathogenesis is hampered by the fact that murine models of infection offer limited resolution for the non-hypervirulent strains which cause the majority of infections. We have performed genome-scale fitness profiling of a multidrug-resistant *K. pneumoniae* ST258 strain during infection of the insect *Galleria mellonella,* with the aim to determine if this model is suitable for large-scale virulence factor discovery in this pathogen. Our results demonstrated a dominant role for surface polysaccharides in infection, with contributions from siderophores, cell envelope proteins, purine biosynthesis genes and additional genes of unknown function. Comparison with a hypervirulent strain, ATCC 43816, revealed substantial overlap in important infection-related genes, as well as additional putative virulence factors that may be specific to ST258. Our analysis also identified a role for the metalloregulatory protein NfeR (also called YqjI) in virulence. Overall, this study offers new insight into the infection fitness landscape of *K. pneumoniae* ST258, and provides a framework for using the highly flexible, scalable *G. mellonella* infection model to dissect the molecular virulence mechanisms of *K. pneumoniae* and other bacterial pathogens.

## Introduction

*Klebsiella pneumoniae* is a gram-negative, capsulated bacterial pathogen responsible for a high proportion of hospital-acquired infections (Podschun and Ullmann 1998; Pendleton, Gorman and Gilmore 2013). Carbapenem-resistant *K. pneumoniae* is classified by the WHO as a critical priority for new drug development (Tacconelli *et al*. 2018). Classical *K. pneumoniae* (c*Kp*) causes a range of opportunistic infections (e.g. pneumonia, skin/soft tissue and catheter-associated urinary tract infections) in the elderly and immunocompromised, while hypervirulent *K. pneumoniae* (hv*Kp*) causes community-acquired invasive disease. Classical and hypervirulent *K. pneumoniae* can be distinguished by the presence of a specific virulence markers, or by their lethality in mice. Though hypervirulent strains are a serious public health threat, the majority of *Klebsiella* disease burden is currently associated with classical strains (Wyres, Lam and Holt 2020). *K. pneumoniae* virulence factors include its protective polysaccharide capsule, O antigen, adhesive pili, capsule overproduction regulators and several different siderophores. Of these, the capsule overproduction genes and the siderophores aerobactin and salmochelin are hypervirulence markers that are absent from the majority of c*Kp* isolates.

Hv*Kp* strains are mouse-virulent with a lethal dose of less than10^6^ colony forming units (cfu), while c*Kp* strains generally are not (Yu *et al*. 2007; Russo and Marr 2019; Russo and MacDonald 2020). The use of mouse models to study c*Kp* infections is typically limited to enumeration of surviving bacteria following challenge with very high inocula (for example, (Diago-Navarro *et al*. 2018; Palacios *et al*. 2018)). Such models capture intermediate points during self-resolving infections and may miss subtle virulence phenotypes. Furthermore, these models cannot easily be used in conjunction with the functional genomics approaches often used to identify infection-related genes *en masse* (e.g. Tn-seq, RNA-seq), as these methods require large numbers of bacteria to provide enough material for sequencing and avoid population bottlenecks (Cain *et al*. 2020). Due to these difficulties the majority of *K. pneumoniae* pathogenesis studies use infection of mice by hv*Kp* strains as the primary measure of virulence. Studying putative c*Kp*-relevant virulence genes in hv*Kp* strains may fail to identify relevant virulence activities, because mechanisms of pathogenesis vary between *K. pneumoniae* isolates (Xiong *et al*. 2015) and because even shared *K. pneumoniae* phenotypes can be underpinned by strain-specific gene sets (Dorman *et al*. 2018; Short *et al*. 2020).

Alternative infection models for *K. pneumoniae* infections include *Dictyostelium discoideum, Drosophila melanogaster, Galleria mellonella, Caenorhabditis elegans,* zebrafish (reviewed in (Bengoechea and Sa Pessoa 2019)), and the *ex vivo* porcine lung (Dumigan *et al*. 2019). Of these models, the greater wax moth *Galleria mellonella* is by far the most established. *G. mellonella* larvae are susceptible to infection by both c*Kp* and hv*Kp* strains (Li *et al*. 2020; Russo and MacDonald 2020), and recapitulate many relevant features of mammalian infections (Insua *et al*. 2013). *G. mellonella* larvae used in research are not standardised, so published lethal doses in this model vary widely (e.g. those of MGH 78578 in (Insua *et al*. 2013; Wand *et al*. 2015), also reviewed in (Pereira *et al*. 2020). Despite this lack of standardisation, the virulence of different *K. pneumoniae* strains in *Galleria* broadly agrees with virulence in mice, though this relationship is not strong enough to reliably differentiate between hv*Kp* and c*Kp* strains in the absence of other information (Li *et al*. 2020; Russo and MacDonald 2020). The flexibility of the *Galleria* model, and its susceptibility to c*Kp* infection, makes this a valuable animal model for high-throughput functional genomics studies of c*Kp*.

We have performed transposon directed insertion sequencing (TraDIS) to identify genes in *K. pneumoniae* RH201207 – a multidrug-resistant c*Kp* strain of the global, outbreak-associated clonal group ST258 – that contribute to *G. mellonella* infection, and compared the *in vivo* fitness requirements of this c*Kp* strain to that of the hv*Kp* strain ATCC 43816. Our results identify known virulence genes along with newly identified putative virulence factors, and we have validated the TraDIS screen with defined single-gene mutants. One gene of interest was the Nickel-dependent transcriptional repressor NfeR, which has not previously been linked to virulence in any species. An NfeR mutant showed reduced virulence and increased expression of a neighbouring ferric reductase gene – effects that were reversed by complementation – but, unexpectedly, could not be linked to any *in vitro* phenotypes. Overall, our results show that the *Galleria mellonella* model is well-suited to high-throughput functional genomics studies of c*Kp* strains, and suggest that even very subtle disruptions to metal homeostasis may be important during c*Kp* infections.

## Material and Methods

### Bacterial strains and culture conditions

Bacterial strains, TraDIS libraries, plasmids and oligonucleotides used in this work are listed in Table S1. *Klebsiella pneumoniae* RH201207 (ST258) and ATCC 43816 (ST493) were grown in LB medium at 37 °C with shaking for routine culture. Where necessary, antibiotics were added in the following concentrations: tetracycline 15 μg mL^-1^, chloramphenicol 25 μg mL^-1^. Viable counts of bacterial cultures were determined by serial dilution in PBS followed by spot-plating of the entire dilution series with technical duplicates.

### *Galleria mellonella* infection experiments

Research-grade *G. mellonella* larvae (Biosystems Technology Ltd, UK) were kept at room temperature in the dark for a maximum of 7 days before use. Injections and haemolymph extractions were performed as described (Harding *et al*. 2013). For survival analyses, *K. pneumoniae* strains were grown to late exponential phase (OD_600_ = 1), harvested by centrifugation, and resuspended in sterile PBS. 10 μL doses of diluted bacterial suspensions were injected into the right hind proleg of the larvae, and the infected larvae were incubated in the dark at 37 °C. Larvae were scored as dead when they were unresponsive to touch.

TraDIS infection experiments were performed in biological triplicate using previously reported high-density mutant libraries. Aliquots of frozen pooled transposon mutant libraries (minimum of 10^8^ cells) were grown overnight, subcultured and grown to late exponential phase, then resuspended and diluted in PBS to an approximate density of 10^7^ cfu mL^-1^. Groups of larvae were injected with 10 μL prepared TraDIS library per larva (approx. dose 10^5^ cfu) and incubated at 37 °C in the dark. Bacteria were extracted from infected *Galleria* as follows: haemolymph was recovered from all of the larvae in each group and pooled in 4 volumes of ice-cold eukaryotic cell lysis buffer (1 × PBS + 1 % Triton X-100) and the mixture was held on ice for ten minutes. Treated haemolymph was centrifuged at 250 × *g* for 5 minutes to pellet eukaryotic cell debris while leaving the majority of bacterial cells in the supernatant. The supernatant was centrifuged at 8000-10000 × *g* for 2 minutes to pellet bacterial cells, and bacteria were resuspended in 5 mL LB and outgrown at 37 °C to approx. OD_600_ of 1 in order to generate enough material for gDNA extraction and sequencing. The final post-infection time points were chosen as the time where there was visible melanisation of the majority of larvae in each group, but no reduction in the volume of recoverable haemolymph. This corresponded to a per-larva bacterial load of ~2 × 10^7^ cfu.

Specific parameters for *G. mellonella* TraDIS infection experiments were as follows. RH201207: 20 larvae per group, 1.5 × 10^5^ cfu inoculum, time points at 2 h post-infection (hpi) and 6 hpi, outgrowth periods 4 hours (2 hpi samples) and 1.5 hours (6 hpi samples); ATCC 43816: 12 larvae per group, 1.1 × 10^5^ cfu inoculum, time point at 4 hpi, outgrowth period 2 hours.

### Genome re-sequencing and annotation of RH201207

Genomic DNA of RH201207 was extracted using the MasterPure Complete DNA and RNA Purification Kit (epicentre) with DNA resuspended in 50 μL nuclease free water by carefully flicking the tube. Purity was checked on a NanoDrop spectrophotometer (260/280 of 2.01, 230/260 of 2.12) and quality and quantity assessed on a TapeStation (Agilent) with DIN of 9.7 and concentration of 36.9 ng μL^-1^.

Nanopore 1D sequencing library was prepared using the genomic DNA by ligation sequencing kit SQK-LSK109 (Oxford Nanopore Technologies, ONT), barcoded using the barcoding extension kit EXP-NPB104 and sequenced on a GridION X5 using a R9.4.1 flow cell (ONT) together with five bacterial genomes from a different study. Bases were called with Albacore v2.0 (ONT) and adapter sequences were trimmed and sequencing reads demultiplexed with Porechop v0.2.3 (https://github.com/rrwick/Porechop). Genome assembly was performed in combination with the previously described paired-end Illumina reads of RH201207 (Jana *et al*. 2017) accessible at the ENA (study PRJEB1730). The hybrid read set was assembled with Unicycler v0.4.7 (Wick *et al*. 2017) using the normal mode and assembly graphs were visualised with Bandage (Wick *et al*. 2015). The final assembly was annotated using Prokka v1.14.5 (Seemann 2014) with additional functional gene annotation by KEGG (Kanehisa and Goto 2000) and UniProt (The UniProt Consortium 2019). Plasmid replicons were identified with PlasmidFinder (Carattoli *et al*. 2014). Typing of the *Klebsiella* K- and O-loci was performed with Kaptive (Wick *et al*. 2018). Iron uptake genes were annotated with SideroScanner (https://github.com/tomdstanton/sideroscanner).

### TraDIS sequencing and analysis

Genomic DNA was extracted by phenol-chloroform extraction. At least 1 μg DNA per sample was prepared for TraDIS as described in (Barquist *et al*. 2016). Sequencing statistics and accession numbers are given in Table S2. Sequencing reads were mapped to the RH201207 or ATCC 43816 genomes using the Bio::TraDIS pipeline as described previously (Langridge *et al*. 2009; Barquist *et al*. 2016), with a 96 % mapping threshold, multiply-mapped reads discarded and one transposon tag mismatch allowed (script parameters: “-v -- smalt_y 0.96 --smalt_r −1 -t TAAGAGACAG -mm −1”). Insertion sites and reads were assigned to genomic features with reads mapping to the 3’ 10 % of the gene ignored, and between-condition comparisons were performed without read count filtering. Gene essentiality was determined by running tradis_essentiality.R (Barquist *et al*. 2016) on the combined gene-wise insertion count data of all three biological replicates. Essential or ambiguous-essential genes of the input samples (the initial transposon library) as defined by the Bio::TraDIS pipeline were excluded from further analysis. Gene functional categories were assigned to genes using EggNOG Mapper (Huerta-Cepas *et al*. 2017) and enrichment of clusters of orthologous groups (COG) was determined by two-tailed Fisher’s exact test in R version 3.6.2 (R-function fisher.test) (R Core Team 2019).

### Mutagenesis and complementation

Deletion mutants of *K. pneumoniae* RH201207 were generated by allelic exchange using vectors derived from pKNG101-Tc as described (Dorman *et al*. 2018). Transposon-insertion mutants of *K. pneumoniae* RH201207 were generated by subjecting the TraDIS mutant library to two rounds of density-gradient selection then identifying mutants by random-primed PCR as described (Short *et al*. 2020).

### *Klebsiella pneumoniae* phenotypic tests

All quantitative phenotypic tests reported are from three biological replicates. Serum susceptibility was determined by incubation of late log-phase cells with 66 % normal human serum (Sigma-Aldrich) as described (Short *et al*. 2020). Siderophore production was visualised by the chrome azurol S assay (Schwyn and Neilands 1987; Louden, Haarmann and Lynne 2011) with the following measures to remove trace iron: glassware was soaked in 6 M HCL for two hours and rinsed three times with ultrapure water, and Casamino acid solution was treated with 27 mL 3 % 8-hydroxyquinoline in chloroform for 20 minutes, then the supernatant was removed, extracted once with a 1:1 volume chloroform, and filter-sterilised. Overnight cultures of *K. pneumoniae* strains were normalised to an OD_600_ of 0.5 prior to spotting on CAS agar, and plates were incubated at 37 °C for 48 hours. For qRT-PCR, total RNA was extracted from bacteria grown in LB medium to OD = 1 using a Qiagen RNeasy kit, and treated with TURBO DNase. Transcripts of *nfeF* and the *recA* housekeeping gene were quantified using a KAPA SYBR fast one-step qRT-PCR master mix according to the manufacturer’s instructions. Three biological replicates and two technical replicates were performed, with 2.5 ng total RNA per reaction. Sensitivity to hydrogen peroxide was determined by diluting stationary phase cultures 1:100 in Mueller-Hinton II medium, then adding hydrogen peroxide (30 % v/v, Sigma-Aldrich) to a final concentration of 4-8 mM. Samples were incubated at 37 °C for 120 minutes before serial dilution and plating to enumerate surviving bacteria. Nickel toxicity and dipyridyl sensitivity tests were performed in a 96-well plate format with 100 μl volume per well and an initial cell density (seeded from overnight culture) of OD_600_ = 0.05. Nickel toxicity tests were performed in LB medium supplemented with nickel(II) sulfate hexahydrate (Sigma-Aldrich). Dipyridyl sensitivity tests were performed in M9 minimal medium supplemented with 0.2 % glucose, with the inoculum washed three times in M9 salts. Plates were sealed with air-permeable film and incubated with shaking for 18 hours at 37 °C prior to measurement of OD_600_.

### Availability of sequencing data

The TraDIS sequencing data is available in the European Nucleotide Archive (https://www.ebi.ac.uk/ena/browser/home) under the Study Accession No. PRJEB20200. Individual sample accession numbers are available in Table S2. Oxford Nanopore reads of RH201207 are available in the ENA repository under study accession number PRJEB40551.

## Results and Discussion

### TraDIS analysis of *G. mellonella* infection determinants in *K. pneumoniae* RH201207

*Klebsiella pneumoniae* RH201207 is a multidrug-resistant isolate of clonal group CG258 used in previous transposon insertion sequencing studies (Jana *et al*. 2017; Short *et al*. 2020). We first measured *K. pneumoniae* RH201207 infection parameters in *G. mellonella*. An inoculum of ~10^5^ cfu was sufficient to kill the majority of infected larvae, with melanisation evident at 5 hours post infection. For TraDIS analysis, three groups of *G. mellonella* larvae were infected with replicate cultures of the *K. pneumoniae* RH201207 mutant pool, and we recovered and sequenced surviving bacteria from the larval haemolymph at 2 hpi and 6 hpi; the volume of haemolymph that could be recovered declined at later time points. Haemolymph was treated with detergent to lyse eukaryotic cells, then bacteria were recovered and grown in rich medium (Fig. 1A). This method allowed high recovery of infecting bacteria without antibiotic selection, while avoiding co-isolation of DNA from either the host or its microbiota. Viable counts were measured on infection and at each sampled time point, and showed approximately seven generations of bacterial replication at 6 hpi (Fig. 1B). Our infection screening parameters therefore allow identification of mutations that impair replication in this host, as well as mutations that cause sensitivity to killing by *G. mellonella* immune system components.

**Figure 1:**
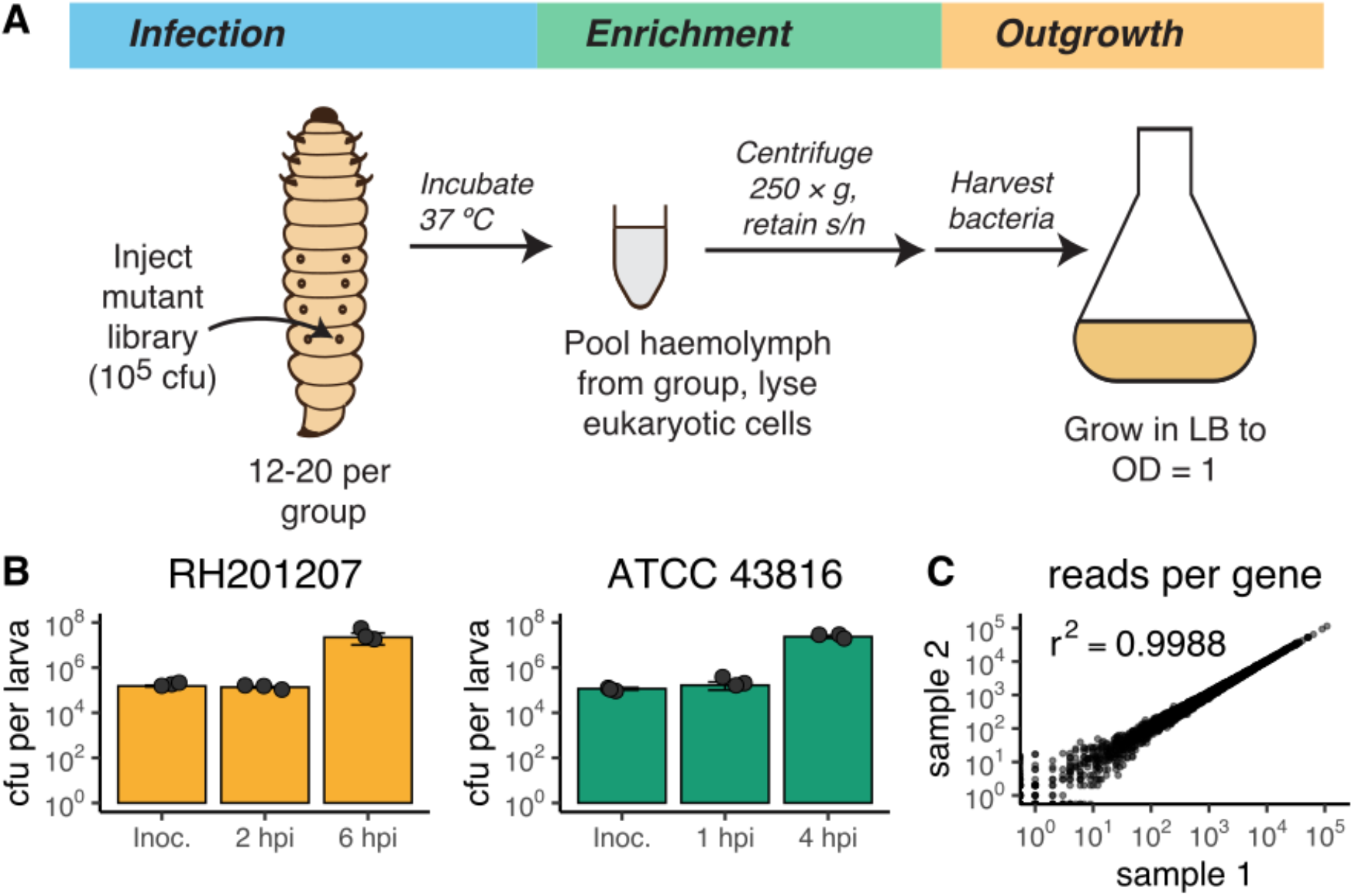
Overview of transposon insertion screening in *Galleria mellonella* larvae. A: Schematic of experimental procedure. Groups of larvae were injected with a pooled library of *Klebsiella pneumoniae* transposon mutants and incubated at 37 °C for up to 6 hours. Haemolymph was extracted from infected larvae, pooled and treated to remove eukaryotic cells. Surviving bacteria were then grown in LB to generate sufficient material for sequencing. B: Per-larva bacterial counts over the course of *G. mellonella infection*. Bacteria were recovered when the load per larva reached approximately 10 million colony forming units (cfu). The experiment was done in triplicates, closed circles represent one measurement, error bars denote standard deviation. Abbreviation: Inoc., inoculum. C: Reads per gene of two biological replicates of the unchallenged RH201207 input transposon library and their Pearson correlation coefficient.

### Re-sequencing of RH201207

During the initial analysis of the TraDIS experiments, we identified a possible mis-assembly of the original RH201207 genome, which could not be improved by optimising the assembly parameters. To improve the assembly and generate a circularised chromosome sequence, we sequenced RH201207 by long-read Oxford Nanopore Technologies (ONT) sequencing.

Nanopore GridION sequencing yielded 2.3 G bases with an average read length of 10.5 kb and maximum read length of 165.9 kb, corresponding to ~400-fold theoretical coverage. Hybrid assembly of these reads together with existing MiSeq reads from Jana *et al* (1.24 million reads, ~35-fold genome coverage) yielded a circularised chromosome of 5,475,789 bp and three circularised potential plasmids (Table 1) of 113,640 bp (contig RH201207_2, IncFIB(pQil) and IncFII(K) replicons), 43,380 bp (contig RH201207_5, IncX3 replicon) and 13,841 bp length (contig RH201207_8, ColRNAI replicon). The remaining contigs of 202,245 bp length could not unambiguously be closed and circularised due to the presence of potential duplicated sequences. Nonetheless, typical plasmid replication proteins and the two replicons IncFIB(K) and IncFII(K) were present on these contigs, indicating that they are likely to be derived from plasmids (Table 1 and Fig. S1). The re-annotated RH201207 genome has 5,787 genes in total (5,546 protein-coding). and encodes a capsule of type KL106 and an O-antigen of type O2v2 as determined by Kaptive (Wick *et al*. 2018).

**Table 1:** Plasmid replicons in RH201207 identified by PlasmidFinder

| Plasmid | Identity | Query length / Template length | Contig | Position in contig | Accession number |
| --- | --- | --- | --- | --- | --- |
| ColRNAI | 100 % | 130 / 130 | RH201207 8 | 10359-10488 | DQ298019 |
| IncFIB(K) | 100 % | 560 / 560 | RH201207 7 | 4026-4585 | JN233704 |
| IncFIB(pQil) | 100 % | 740 / 740 | RH201207 2 | 217-956 | JN233705 |
| IncFII(K) | 97.8 % | 148 / 148 | RH201207 2 | 50275-50422 | CP000648 |
| IncFII(K) | 97.8 % | 148 / 148 | RH201207 6 | 1728-1875 | CP000648 |
| IncX3 | 100 % | 374 / 374 | RH201207 5 | 32841-33214 | JN247852 |

### Identification of infection-related genes

TraDIS sequencing, read mapping and quantification of each transposon insertion site was performed using the Bio::TraDIS pipeline (Barquist *et al*. 2016). Each sample yielded from 12.9 million to 15.1 million transposon-tagged reads, > 89 % of which unambiguously mapped to the RH201207 chromosome (excluding unscaffolded contigs and plasmids) (Table S2). Analysis of the unchallenged RH201207 TraDIS library showed a total of more than 500,100 unique transposon insertion sites distributed across the chromosome, which corresponds to one insertion in every eleven nucleotides (Table 2). 638 genes (or 11.84 % of the 5390 chromosomal genes) were either essential or ambiguous-essential as defined by the Bio::TraDIS pipeline (Barquist *et al*. 2016), which is in good concordance with the first description of this library (Jana *et al*. 2017). A challenge in applying highly saturated transposon insertion libraries to infection models is the occurrence of bottlenecks, that is a stochastic drastic reduction in population size, which results in reduced genetic diversity of the population (Cain *et al*. 2020). We found a slight reduction in the diversity of the mutant library post-infection, with recovered unique insertion sites of nearly 380,000 and 360,000 total insertions (of 500,100) at 2 hpi and 6 hpi, respectively, however transposon insertion density remained very high with an insertion approximately every 15 bp (Table 2). To test if the loss of mutant library diversity had compromised the resolution of our experiment, we compared individual replicates using linear correlation analyses. Pearson correlation coefficients between *in vivo* replicates were very high with *r*^2^ values greater than 0.98 when analysing reads per gene and insertion indices per gene, and 0.69 to 0.77 when analysing the reads per unique insertion site (Fig. 1, S2, S3 and Table S3). Therefore, although our TraDIS experiments show a mild bottleneck, this is highly unlikely to affect any downstream analyses that use gene-level metrics. To identify *in vivo* fitness genes, the numbers of transposon insertion reads within each gene were compared between the *Galleria* infection and the inoculum pools, with essential and ambiguous-essential genes excluded to reduce false positives. Our analysis identified mutants of 133 (of 4,752) nonessential genes to be significantly less abundant at 6 hpi and two features, *rseA* and KPNRH_05271 to be slightly, but significantly enriched (Fig. 2 and supplementary table S4). 35 genes were so severely depleted after infection that they can be considered conditionally essential in *G. mellonella* (Table S4).

**Table 2:** Unique insertion sites of the TraDIS libraries used

| Sample | Chromosome length | Unique Insertion Sites (UIS) | Length / UIS |
| --- | --- | --- | --- |
| RH201207 | 5,475,790 bp | 510,834 | 10.72 bp |
| RH201207 2 hpi | 5,475,790 bp | 379,343 | 14.43 bp |
| RH201207 6 hpi | 5,475,790 bp | 359,602 | 15.23 bp |
| ATCC 43816 | 5,374,835 bp | 415,608 | 12.93 bp |
| ATCC 43816 4 hpi | 5,374,835 bp | 350,363 | 15.34 bp |

**Figure 2:**
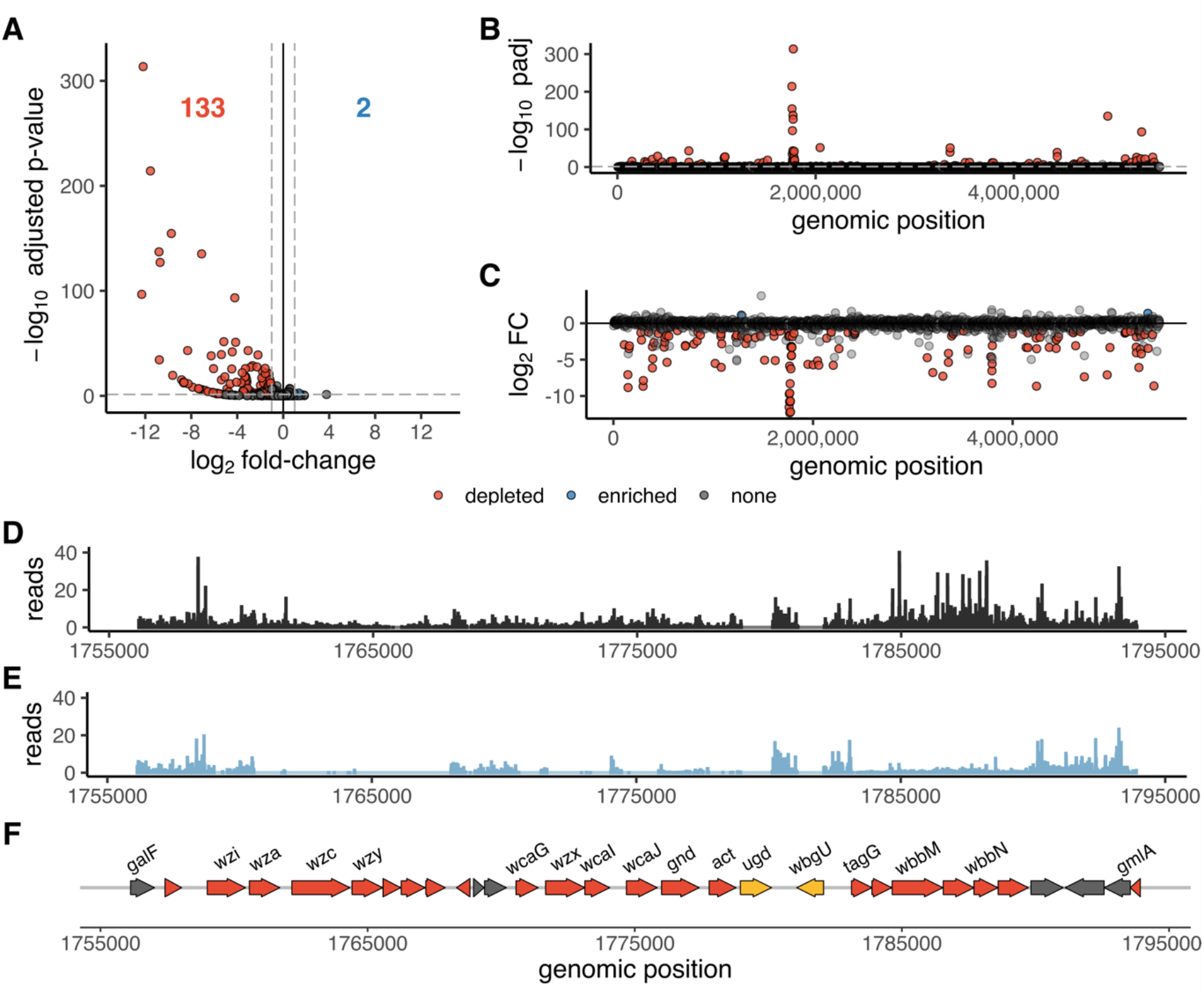
TraDIS analysis identifies multiple potential *K. pneumoniae* fitness factors important during *G. mellonella* infections. A: Volcano plot showing significantly less abundant genes 6 h after infecting *G. mellonella* larvae in red and significantly higher abundant genes in blue. Non-significant genes are shown in grey. Genes were considered significant if they possessed a Benjamini-Hochberg corrected *p*-value below a threshold of 0.05 and an absolute log2 fold-change greater than 1. Essential and ambiguous-essential genes are removed from the analysis. B and C: Manhattan plots with Benjamini-Hochberg corrected *p*-value and log2 fold-change on the y-axis, respectively. Abbreviations: padj, Benjamini-Hochberg corrected *p*-value; FC, fold-change. D and E: Reads per nucleotide of the inoculum and at 6 hpi, respectively of the chromosomal region containing the K- and O-locus. For this graphical representation, triplicate TraDIS samples were combined and reads were normalised to the total number of reads in all three replicates. F: Gene annotations of the *Klebsiella* K-locus (*galF* to *wbgU*) and O-locus (*tagG* to *gmlA*), encoding the polysaccharide capsule and lipopolysaccharide O antigen, respectively are coloured according to the TraDIS results at 6 h post-infection; significantly depleted genes are shown in red, essential genes which were not included in the analysis in yellow and genes without significant changes in grey.

### Surface polysaccharides, cell envelope and iron acquisition genes are critical to c*Kp G. mellonella* infection

#### Capsule, lipopolysaccharide and enterobacterial common antigen

Capsule and lipopolysaccharide (LPS) are essential to the virulence of *K. pneumoniae*, (Podschun and Ullmann 1998). The capsule allows *K. pneumoniae* to circumvent host detection and prevent an early immune response; acapsular mutants are less virulent in mouse models and are unable to spread systemically (Paczosa and Mecsas 2016). The lipopolysaccharide, consisting of lipid A, core and O antigen is able to bind and sequester parts of the complement system (Paczosa and Mecsas 2016). Both surface polysaccharides also play a role in modulating innate immunity (Bengoechea and Sa Pessoa 2019). Following *G. mellonella* infection, the eight genes with the most dramatic reduction in read counts (log2 fold-change < −12) all belonged to the capsule or O-antigen loci (Fig. 2B & C), and almost all genes in these clusters were significantly depleted (Fig. 2D-F).

Four genes (*wzxE, wecD, rffH, wecA*) of the Enterobacterial common antigen (ECA) synthesis locus (encoded by genes *rho*/KPNRH_05213 to yifK/KPNRH_05200) were also significantly depleted at 6 hpi (Fig. 2B & C). The ECA is a conserved carbohydrate antigen common to most Enterobacterales and plays an important role in bacterial physiology and its interaction with the environment (Rai and Mitchell 2020). *K. pneumoniae* ECA synthesis mutants were attenuated in *in vitro* murine lung and spleen tissues infections, but not *in vivo* in a murine intranasal infection model (Lawlor *et al*. 2005). Therefore, our analysis identified mutants of all three major polysaccharide antigens (O, K and ECA) present on the cell surface (Rai and Mitchell 2020) to be attenuated in *G. mellonella*.

#### Cell envelope proteins and regulators

Genes that encode cell envelope proteins appeared to be important during *G. mellonella* infection. The major outer membrane lipoprotein Lpp (also known as Braun’s lipoprotein) encoded by *lpp*/KPNRH_02211, and the genes of the Tol-Pal system (*pal/*KPNRH_03757, *tolB*/KPNRH_03758, *tolA*/KPNRH_03759 and tolQ/KPNRH_03761) (located at 3.8 Mbp) were significantly depleted after 6 hpi. The Tol-Pal system is involved in maintaining outer membrane integrity and consists of five proteins: TolA, TolQ, and TolR in the inner membrane, TolB in the periplasm, and the peptidoglycan-associated lipoprotein Pal anchored to the outer membrane (Lloubès *et al*. 2001). Lpp is a crucial protein in the outer membrane covalently linking it with the peptidoglycan layer (Asmar and Collet 2018) and it is important for complement resistance in multiple, phylogenetically distinct *K. pneumoniae* strains (Short *et al*. 2020). Mutations in both Lpp and Tol–Pal have been previously linked to attenuated virulence in diverse bacteria (Sha *et al*. 2008; Godlewska *et al*. 2009; Asmar and Collet 2018). One of the most abundant proteins in the outer membrane is the porin OmpK36 (Hernández-Allés *et al*. 1999) and mutants thereof were attenuated in a *G. mellonella* model (Insua et al. 2013) and a pneumonia mouse model (March et al. 2013). Likewise, our experiments identified a significant underrepresentation of transposon insertion mutants in *ompK36* (KPNRH_01613) 6 hours post-infection.

Also required for infection were regulators of cell envelope composition and integrity. Mutants of *phoPQ* were significantly less abundant following infection. PhoP-PhoQ is a two-component system, comprising the inner membrane sensor PhoQ and the cytoplasmic regulator PhoP. This system regulates lipid A remodelling in *K. pneumoniae in vivo* and *in vitro* (Llobet *et al*. 2015) and other virulence-associated genes in many enteric pathogens (Groisman 2001; Bijlsma and Groisman 2005; Alteri *et al*. 2011; Lin *et al*. 2018). *Salmonella* Typhimurium knockout strains of*phoP* and *phoQ* are highly attenuated for virulence in macrophages and a mouse infection (Miller, Kukral and Mekalanos 1989) and this two-component system also makes a small contribution to *K. pneumoniae* virulence during *G. mellonella* infections (Insua *et al*. 2013). We also noted significant changes in mutant abundance of genes related to the alternative sigma-factor RpoE (σ^24^ or σ^E^), one of the major regulators of cell envelope stress response systems (Flores-Kim and Darwin 2015; Roncarati and Scarlato 2017) (Treviño-Quintanilla, Freyre-González and Martínez-Flores 2013). RpoE activity is tightly controlled by a proteolytic cascade: after dissociation of RseB from RseA (the anti-sigma factor which binds RpoE), RseA is partially cleaved by the proteases DegS and RseP, then fully degraded by other cellular proteases such as ClpP/X-A, Lon and HslUV, releasing RpoE in the cytoplasm (Roncarati and Scarlato 2017). While *rpoE* itself and *rseP* are essential genes and therefore not included in our analysis, transposon insertion mutants of *degS, clpP, clpX* and *lon* were significantly less abundant, highlighting the role of RpoE gene regulation during infection. Transposon mutants of the RpoE inhibitor RseA and insertions in the promoter of the RseA-degrading protease HslUV (KPNRH_05271) were enriched after infection, indicating that increased RpoE signalling can enhance fitness in *G. mellonella*. Both RseA and HslUV are, via RpoE, involved in stress response and the regulation of virulence genes in multiple Enterobacteriaceae species (Flores-Kim and Darwin 2015). Interestingly, none of the RpoE regulated cell envelope stress response systems CpxAR, BaeRS, Rcs, and Psp (Flores-Kim and Darwin 2015) were identified in our screen, indicating either a redundancy in these systems or the involvement of other factors. Such factors could be the periplasmic chaperones Skp and SurA; both are members of the RpoE regulon in *E. coli* (Dartigalongue, Missiakas and Raina 2001). Mutants of *skp* and *surA* were less abundant in *Galleria* TraDIS and have previously been shown to be involved in pathogenicity in *E. coli*, *Salmonella* Typhimurium, *Shigella flexneri* and *Pseudomonas aeruginosa* (Sydenham *et al*. 2000; Redford and Welch 2006; Purdy, Fisher and Payne 2007; Klein *et al*. 2019).

#### Iron acquisition systems

Iron acquisition is essential during *K. pneumoniae* infections, and RH201207 produces the siderophores enterobactin and yersiniabactin to chelate host iron. Because siderophores confer benefits to bacterial populations, rather than individual cells, loss of siderophore biosynthesis genes is not expected to influence fitness in a mutant pool, and siderophore synthesis genes were not identified by the *Galleria* infection TraDIS. Utilisation of enterobactin was, however, important: mutations in three of the four ferric iron-enterobactin uptake complex components *tonB* (KPNRH_02155), *exbB* (KPNRH_00722), and *exbD* (KPNRH_00723) reduced fitness in *G. mellonella*. TonB is essential for pathogenicity in multiple bacteria, including hypervirulent *K. pneumoniae* (Hsieh *et al*. 2008). The fourth component of this complex, *fepB*, was present in multiple copies in the RH201207 genome, so presumably mutation of just one gene does not cause loss of function. Mutation of the yersiniabactin receptor gene *fyuA* did not impair fitness, suggesting that enterobactin is sufficient to allow bacterial replication in the *G. mellonella* haemolymph. This is consistent with the roles of *K. pneumoniae* siderophores in murine infections, where enterobactin alone allows replication in serum by sequestering iron from transferrin (Bachman *et al*. 2012). Transferrin is also present in the haemolymph of *G. mellonella* (Vogel *et al*. 2011).

#### Further genes involved in *Galleria mellonella* fitness

Additional metabolic and hypothetical genes also contributed to fitness during *Galleria* infection. These included the majority of genes in the *aro* operon for synthesis of chorismite (a precursor to aromatic amino acids), *cys* genes for sulphate assimilation and cysteine biosynthesis, purine biosynthesis genes, several components of the electron transport complex, and *sspAB* stringent response proteins. Only 11 genes of unknown function were identified as infection-related (note some K- and O-locus genes were initially annotated as hypothetical). Several transcription regulators were also identified that compromised fitness when mutated: these were the arginine/lysine synthesis repressor *argP*, the fatty acid regulator *fabR*, the zinc-dependent repressor *nrdR* and the nickel-dependent repressor *yqjI*.

#### Temporal analysis of the *K. pneumoniae* RH201207 *G. mellonella* infection

We also analysed bacteria at 2 hpi, in order to gain insight into the events of early infection, and distinguish mutations influencing survival in the presence of *G. mellonella* immune system components from those influencing replication in this host. Mutants in 54 genes were significantly less abundant at 2 hpi, and only mutants lacking the cell division protein FtsB were enriched (Fig. 3, S4-S6 and supplementary table S4). Comparison of both timepoints showed 18 genes depleted only at 2 hpi, 36 genes less abundant at both timepoints and 97 genes for which mutant abundance was reduced only at 6 h post-infection (Fig. 3A). The genes that were depleted at both time points consists of two subsets: 25 genes that are depleted within the first 2 h and whose abundance does not change further, and 11 genes that are depleted at 2 hpi and then further decrease in abundance over time. Ten out of these 11 genes are part of the K-locus, the other one is *tagG*, the first gene of the neighbouring O antigen locus.

**Figure 3:**
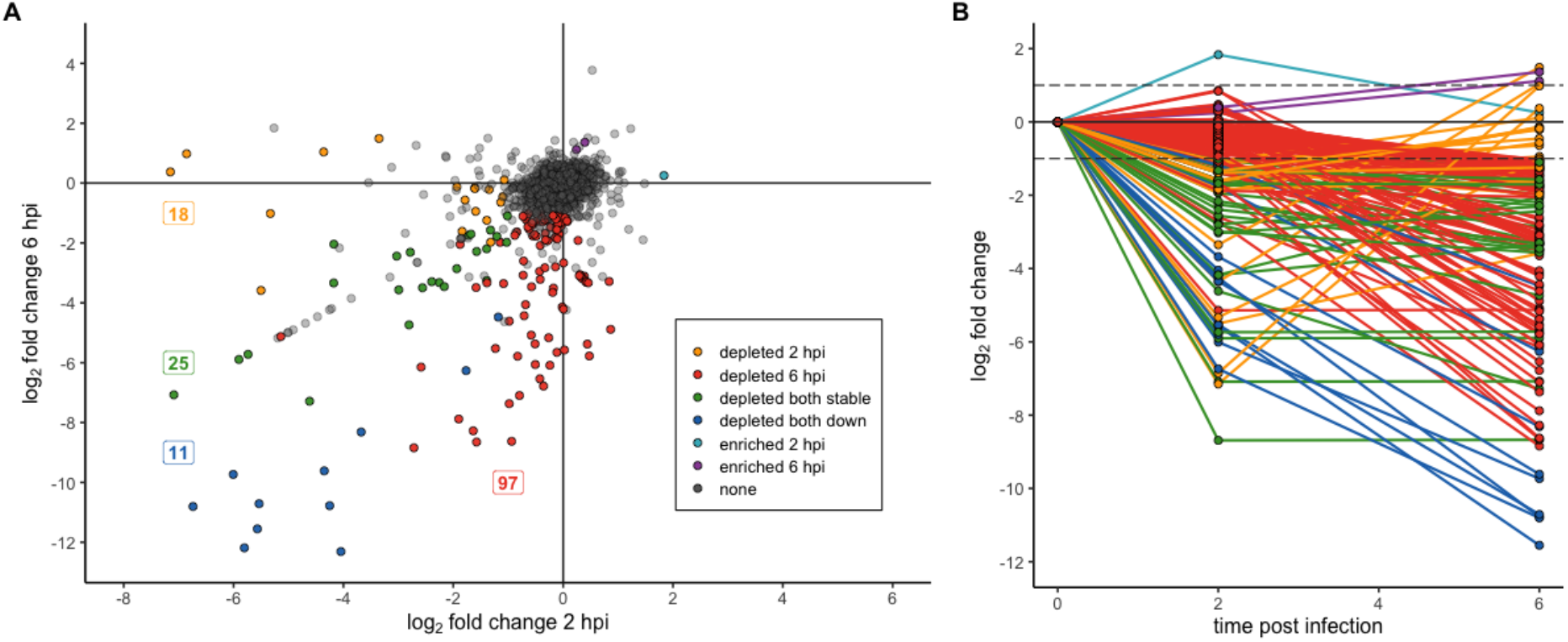
Few genes are important for the onset but not the late stages of a *G. mellonella* infection. A: The plot shows log2 fold-changes at 2 hpi and 6 hpi on the x- and y-axis, respectively. Transposon mutants significantly less abundant at 2 hpi and 6 hpi are shown in yellow and red, respectively. Mutants which are depleted at both timepoints are shown in either green or blue (if the log2 fold-change at 2 hpi roughly equals that at 6 hpi in green and if their abundance was much lower at 6 hpi in blue). Genes significantly higher abundant at 2 hpi and 6 hpi are shown in turquois and purple, respectively. Non-significant genes are shown in grey. Numbers indicate the number of genes of each group. B: Graph highlighting the direction of the log2 fold-change change over time. Genes are coloured according to A; non-significant genes are not shown.

Of the 18 genes less abundant only at the beginning of the infection, most were barely within our threshold for fitness-related genes. But six genes had log2 fold-changes of −3 up to −7, among them two tRNAs, one hypothetical protein and the three genes *oxyR* (KPNRH_05248), *rnhA* (KPNRH_04380) and *tadA* (KPNRH_01302). It is very likely that the tRNAs are false positives due to their very short length and therefore low number of insertions. OxyR is a conserved LysR-type transcription factor which plays a key role in the regulation of defence mechanisms against oxidative stress (Christman, Storz and Ames 1989), and has previously been linked to *K. pneumoniae* pathogenesis (Hennequin and Forestier 2009). RnhA is the ribonuclease HI that cleaves RNA of RNA–DNA hybrids and is involved in DNA replication, DNA repair, and RNA transcription (Kochiwa, Tomita and Kanai 2007) and *tadA* encodes a tRNA-specific adenosine deaminase. This protein is responsible for adenosine to inosine RNA editing of tRNAs and mRNAs, and is involved in the regulation of a toxin-antitoxin system in *E. coli* (Bar-Yaacov *et al*. 2017).

Of all significantly less abundant insertion mutants, the by far largest set, consisting of 97 genes, is depleted only at the later timepoint 6 h after infection. This group of genes contains for example the two-component system *phoPQ*, the Tol–Pal system and the majority of genes of the O antigen cluster. Our data indicates that these genes might only have a minor impact on fitness during very early stages of infection.

### Shared *in vivo* fitness determinants in c*Kp* RH201207 and hv*Kp* ATCC 43816

We then sought to compare the *G. mellonella* fitness landscape of the c*Kp* ST258 strain RH201207 with a representative hv*Kp* strain. *G. mellonella* infection TraDIS was performed with the well-studied mouse-virulent strain ATCC 43816, which is an ST493 strain possessing type O1v1 O antigen and K2 capsule type. Experiments were performed in the same way as for RH201207 except that bacteria were recovered for sequencing at 4 hpi, due to the faster progression of infection when using this strain (Fig. 1B). Each sample of ATCC 43816 yielded from 1.85 million to 2.08 million reads, more than 96 % of which were reliably mapped to the chromosome of ATCC 43816 KPPR1 (CP009208.1) (Table S2). The insertion density of the unchallenged ATCC 43816 mutant library was 415,000 or one insertion per 13 nucleotides, which is similar to the RH201207 library (Table 2, S2), and 502 genes were classified as essential or ambiguous-essential (9.62 % of 5217 genes). Reproducibility between experimental replicates was very high in this experiment, with Pearson correlation coefficient > 0.97 for reads or insertion indices per gene, and > 0.91 when comparing reads per unique insertion site (Fig. S2, S3 and Table S3).

We identified 92 nonessential genes of *K. pneumoniae* ATCC 43816 that had significantly lower mutant abundance after the *Galleria* infection and two genes (VK055_RS06420 and VK055_RS19345) that were enriched (Fig. S4-S6 and supplementary table S5). Mutants in 10 genes of the 18-gene K locus of ATCC 43816 were significantly depleted after *G. mellonella* infection, thereby mirroring the results of RH201207. In contrast, genes of the O-locus were not implicated in *in vivo* fitness in *G. mellonella*. This was unexpected because our previous study showed that O antigen genes were required for ATCC 43816 serum resistance (Short *et al*. 2020) and we predicted that resistance to *G. mellonella* humoral immunity may have similar requirements. However other studies of ATCC 43816 O antigen have shown variable effects on virulence and related *in vitro* phenotypes (Shankar-Sinha *et al*. 2004; Yeh *et al*. 2016). We speculate that the ATCC 43816 capsule has a dominant role in protection from *G. mellonella* immunity and masks the activity of O antigen.

To compare the global TraDIS results from RH21207 with those of ATCC 43816, we identified their shared genes by bidirectional best hits BLAST search using a sequence similarity cut-off of 80 %. This analysis identified 4,391 shared genes, 3,974 of which were non-essential in both strains and therefore included in the comparison. We identified 149 shared genes with a role in *G. mellonella* infection in either RH201207 or ATCC 43816, and 44 genes were implicated in both strains (Fig. 4A). This means that 37.3 % of the hits in RH201207 were also identified in ATCC 43816 and 58.7 % of all hits in ATCC 43816 were identified in either of the two RH201207 datasets, the largest overlap was with the later timepoint of 6 hpi. Amongst those genes were, for example, the membrane-associated genes *ompK36, tolABQ, and lpp*, the outer membrane protein assembly factor *bamB/yfgL*, the periplasmic chaperone *surA*, and the ECA synthesis genes *rffG* and *wecA*.

**Figure 4:**
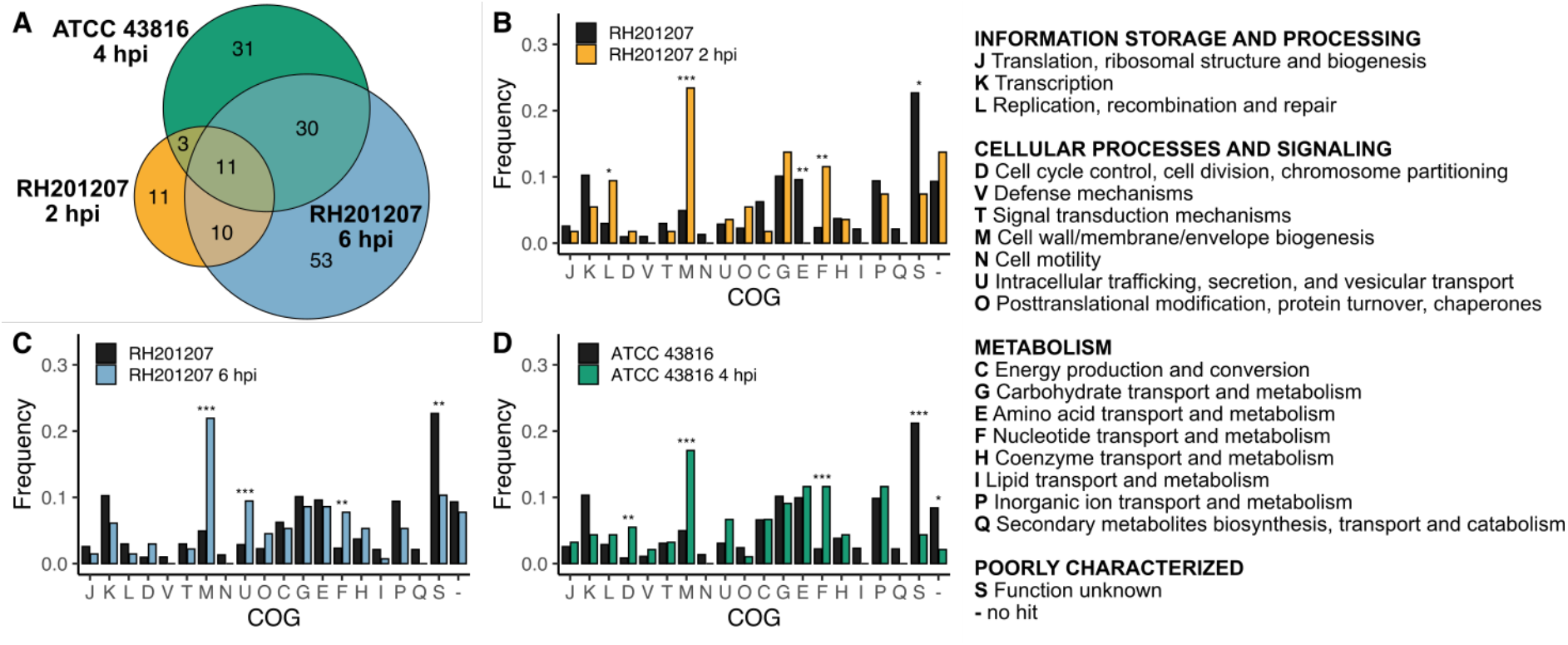
Multiple genes, including the cps cluster contribute to the fitness during *Galleria* infections of both classical and hypervirulent *K. pneumoniae*. A: Venn diagram showing the overlap of significantly less abundant insertion mutants in the classical ST258 strain RH201207 and the hypervirulent strain ATCC 43816. Only non-essential genes shared by both strains as determined by a bi-directional best blast analysis are shown. B-D: Cluster of orthologous groups (COG) enrichment analysis shows an overrepresentation of outer membrane biogenesis as well as nucleotide transport and metabolism genes in all sets of significantly depleted genes during *G. mellonella* infection. Genes were assigned to Cluster of Orthologous Groups (COG) (Tatusov *et al*. 2000) with eggNOG. The bars represent the percentage of genes that belong in that category. Black bars denote the frequency of COGs in all non-essential genes of the particular strain, the frequency of all significantly depleted genes after *Galleria* infection is coloured as follows, A: RH201207 2 hpi, B: RH201207 6 hpi, C ATCC 43816 4 hpi. *P*-values were determined by Fisher’s exact test in R and the level of significance is indicated by asterisks (*, *P* <0.05; **, *P* < 0.01; ***, *P* < 0.001). COGs without genes assigned to them (A, B, R, W, Y & Z) were removed.

We performed a clusters of orthologous groups (COG) enrichment analysis to define and compare the broad pathways and molecular mechanisms that may contribute to fitness during *G. mellonella* infections in both strains. The COG classifications of the genes in the RH201207 and the ATCC 43816 genome was annotated with EggNOG Mapper (Huerta-Cepas *et al*. 2017) and the COGs of all genes in which mutants were significantly underrepresented after infection were compared to the total gene set. In RH201207, genes of the replication, recombination and repair (L), cell wall/membrane/envelope metabolism (M) and nucleotide transport and metabolism (F) clusters were overrepresented among the infection-related genes at 2 hpi, whereas amino acid transport and metabolism (E) and genes of unknown function (S) were underrepresented (Fig. 4B). At 6 h post-infection, cell wall/membrane/envelope metabolism (M), intracellular trafficking, secretion, and vesicular transport (U) and nucleotide transport and metabolism (F) were overrepresented, whereas only genes of unknown function (S) were underrepresented (Fig. 4C). In the ATCC 43816 infection determinants, genes assigned to carbohydrate transport and metabolism (G), cell wall/membrane/envelope biogenesis (M) and nucleotide transport and metabolism (F) were overrepresented. The cell wall/membrane/envelope metabolism (M) and nucleotide transport and metabolism (F) clusters were the only ones in which infection-related genes were overrepresented for both c*Kp* and hv*Kp*, and at early and late time points. This finding further stresses the importance of cell membrane integrity and surface polysaccharides in infections, but also demonstrates the essentiality of nucleotide metabolism during the course of an infection. The latter COG included the genes *purA, purC, purE, purH*, and *rdgB;* all purine biosynthesis genes that showed significantly reduced fitness in both strains. The ability to *de novo* synthesize purines has been associated with the intracellular survival of bacterial pathogens such as *Burkholderia pseudomallei, Shigella flexneri, and* uropathogenic *Escherichia coli* (Ray *et al*. 2009; Shaffer *et al*. 2017).

### Comparison with genetic fitness requirements in murine hosts and *in vitro* virulence screens

We then sought to compare our findings in *G. mellonella* to published results of other high-throughput fitness screens of *K. pneumoniae*, with the caveat that such screens by their nature provide only a snapshot of the events of an infection, and are unlikely to be comprehensive. The *G. mellonella* fitness genes identified in both strains showed considerable similarity to those required for survival in human serum, which we examined previously using the same mutant libraries (Short *et al*. 2020). This was largely due to the importance of capsule, O antigen and cell envelope proteins such as Lpp for survival under both selections; the many metabolic genes identified as infection-relevant, for example those of purine biosynthesis (Fig. 4), generally did not contribute to serum survival. For RH201207, over half of the genes identified as required for full serum fitness also contributed to fitness in *G. mellonella*.

Fitness factors required for intestinal colonisation of mice have also been examined in an ST258 background (Benoit *et al*. 2019), although this screen was not comprehensive due to various experimental factors. Several genes required in *G. mellonella* are also required for intestinal colonisation, such as *bamB* (an outer membrane assembly protein), *ompC/ompK36, cyaA*, (adenylate cyclase) and *typA* (a GTP-binding protein). Finally, there are some important common factors among the requirements for infection of *G. mellonella*, and for an hv*Kp* murine lung infection (Paczosa *et al*. 2020). Genes contributing to both infection types include some of those encoding ubiquitous virulence factors such as capsule or siderophore importers. Notably, genes involved in aromatic amino acid biosynthesis (e.g. *pabAB* aminodeoxychorismate synthase, *aro* operon genes), and purine biosynthesis (e.g. *purH)* were required both for murine lung infection, and for *Galleria mellonella* infection in both strain backgrounds. The importance of these pathways for two very different types of infection, in representatives of both hv*Kp* and c*Kp*, suggests that they may be general infection requirements for *K. pneumoniae*.

### Validation of transposon insertion sequencing results and investigation of NfeR activity

Five single-gene mutants of *K. pneumoniae* RH201207 were tested for lethality in *G. mellonella* to validate the TraDIS screen. The validation set included transposon insertion mutants in the transcription antiterminator *rfaH*, the capsule gene *wzc* and the enterobacterial common antigen gene *wzxE*, as well as clean deletion mutants of*phoQ* and *nfeR*. Researchgrade *G. mellonella* larvae were injected with 10^6^ cfu of each strain, or a PBS control, and monitored for up to 72 hours. All strains showed a statistically significant virulence defect (Fig. 5A), with the exception of RH201207 Δ*phoQ*; larvae infected with the Δ*phoQ* mutant showed increased survival but the degree did not reach statistical significance. It is possible that the TraDIS screen detected changes that reflect fitness in a competition environment with other strains, or that are important at early stages of infection but are not relevant in the longer term.

**Figure 5:**
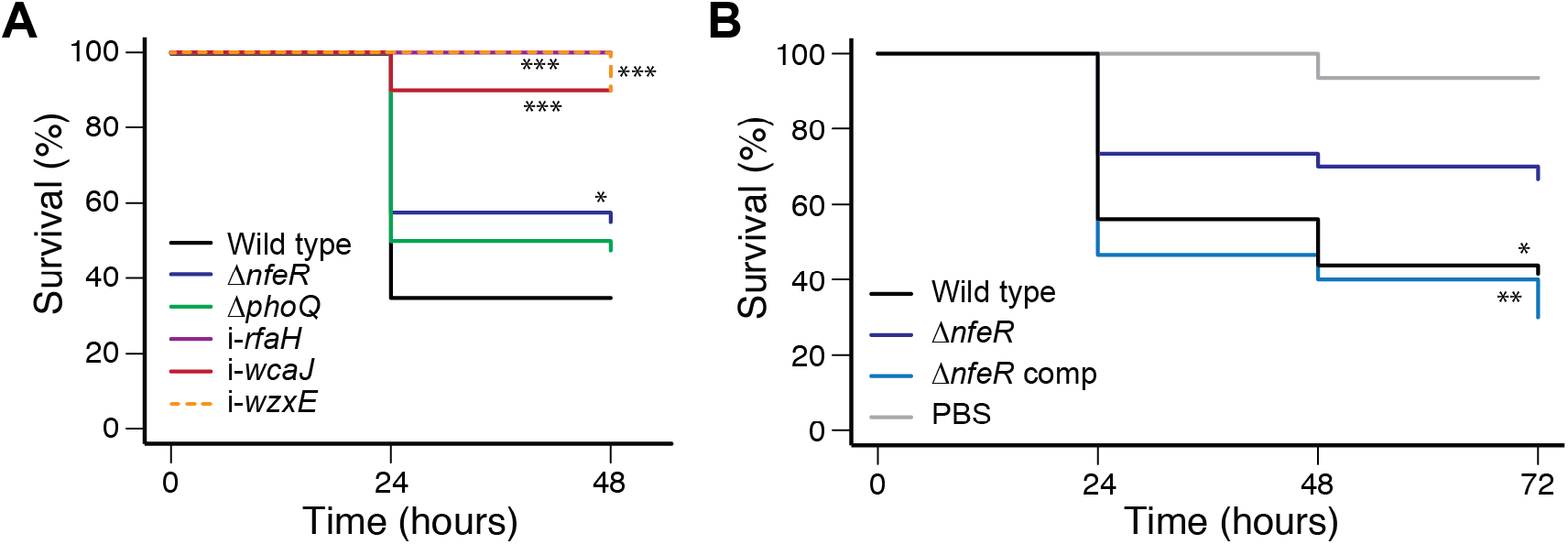
Validation of TraDIS results with single-gene knockouts. A: Survival curves for *G. mellonella* larvae following infection with *K. pneumoniae* RH201207 and mutants in defined genes. B: Complementation of the virulence defect of RH201207 Δ*nfeR*. Mutants where survival is significantly different to wild-type (Kaplan-Meier test) are indicated by asterisks (*, *P* <0.05; **, *P* < 0.01; ***, *P* < 0.001).

We selected *nfeR* (KPNRH_00645 also called *yqjI*) for further characterisation, as this gene has not previously been linked to virulence in any species. In our TraDIS experiments, this gene was required in *K. pneumoniae* RH201207, but not in ATCC 43816. Complementation of the Δ*nfeR* mutation restored the virulence of the wild-type RH201207 strain (Fig. 5B). NfeR contributes to metal homeostasis in *E. coli* by repressing expression of a neighbouring ferric reductase gene, *nfeF* (Fig. 6A); this repression is relieved under excess nickel conditions (Wang, Wu and Outten 2011; Wang *et al*. 2014). High nickel levels can disrupt iron homeostasis in *E. coli*, leading to a longer lag phase due to reduced accumulation of iron (Rolfe *et al*. 2012; Washington-Hughes *et al*. 2019). We hypothesised that loss of *nfeR* may lead to deregulated metal homeostasis in *K. pneumoniae* RH201207, resulting in reduced virulence in *Galleria mellonella*.

**Figure 6:**
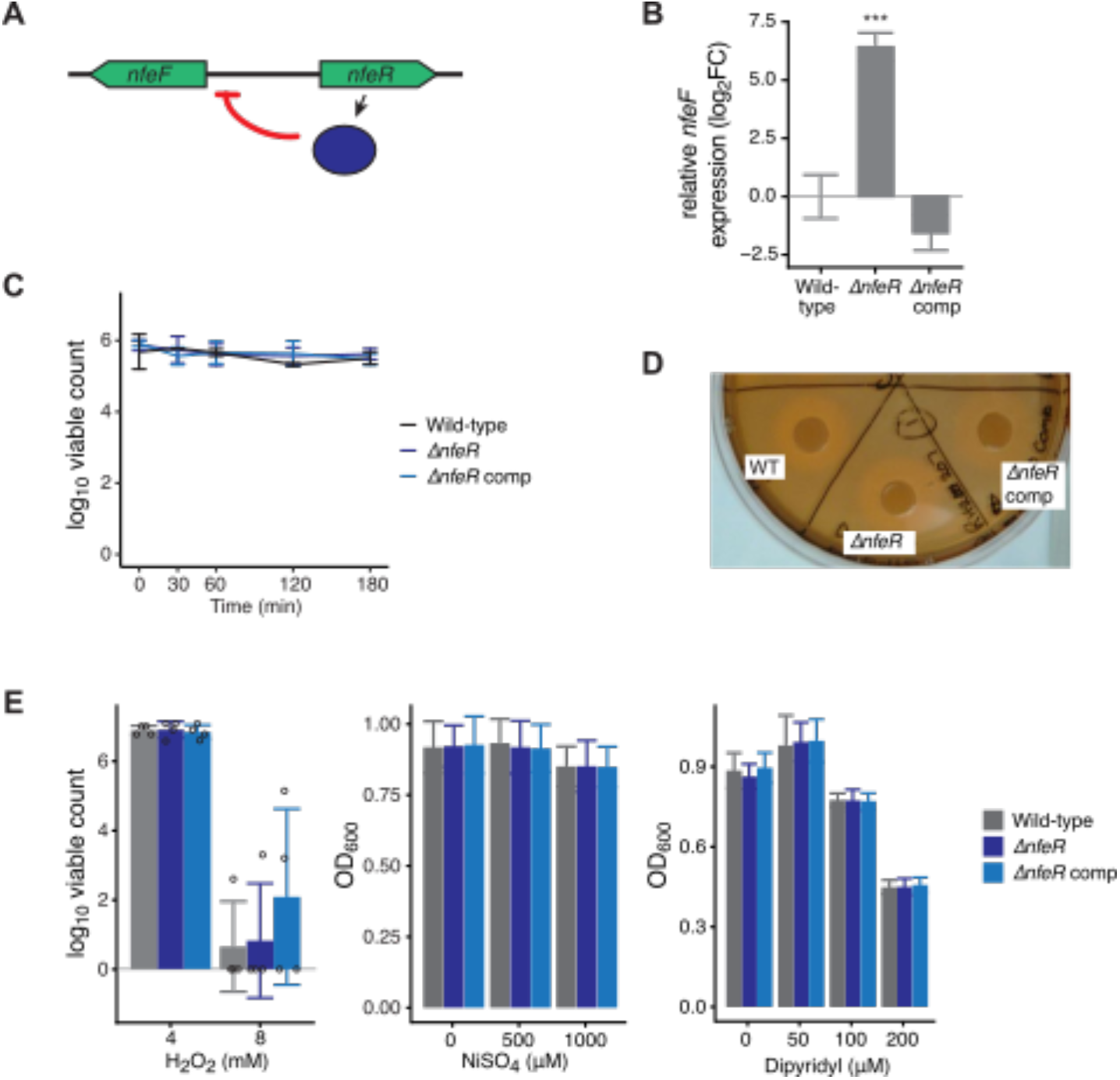
Investigation of NfeR function in RH201207. Mutation of *nfeR* resulted in a dramatic increase in *nfeF* expression. A: Schematic of *nfeF* gene expression regulation by NfeR. B: Deletion of *nfeR* leads to significant overexpression of *nfeF* as measured by qRT-PCR. Level of significance in comparison to RH201207 is indicated by asterisks (***, *P* < 0.001). C: Serum survival of RH201207 Δ*nfeR*. Late exponential phase bacteria were incubated in 66 % normal human serum for 3 hours, and viable counts measured over time. D: Siderophore production by RH201207 Δ*nfeR*, using chrome azurol S agar assay. Results shown are representative of two independent experiments, each comprising at least two biological replicates. E: Sensitivity of RH201207 Δ*nfeR* to hydrogen peroxide, Nickel toxicity and dipyridyl-induced metal starvation. Stationary phase cells were diluted directly into H_2_O_2-_ supplemented Mueller Hinton broth, and surviving cells enumerated after 100 min. For Nickel and dipyridyl toxicity, growth was measured after incubation for 18 hours at 37 °C. Results shown are the mean and standard deviation of four (H_2_O_2_) or three biological replicates, which in the case of Ni and dipyridyl comprised of 3 technical replicates.

We first tested whether NfeR regulates *nfeF* expression in *K. pneumoniae* RH201207 by qRT-PCR using RNA extracted from late exponential phase bacteria. As shown (Fig. 6B), deletion of *nfeR* caused a dramatic 85-fold increase in *nfeF* expression, and this change was not seen in the complemented strain. We hypothesised that the virulence defect of RH201207 Δ*nfeR* may arise from a reduced defence against humoral immunity, or reduced ability to acquire iron, as *G. mellonella* recapitulates several relevant features of mammalian serum-based immunity, and is iron-limited (Lucidi *et al*. 2019). However, the mutant did not show any differences relative to its wild type either in its ability to withstand serum exposure (Fig. 6C), or its siderophore production (Fig. 6D). Finally, we examined three phenotypes that may be disrupted when metal homeostasis is altered: resistance to oxidative stress (hydrogen peroxide), nickel toxicity and sensitivity to iron starvation. Sensitivity to oxidative stress is influenced by cellular iron pools, and was previously shown to be growth-phase dependent for this reason (Touati 2000). Homologues of *nfeR* have been implicated in resistance to nickel toxicity in large-scale fitness screens (Price *et al*. 2018). It was also shown that some genes in the same pathway as *nfeF* are more sensitive to dipyridyl-mediated iron starvation (McHugh *et al*. 2003). However, *K. pneumoniae* RH201207 Δ*nfeR* did not show changes in any of these phenotypes. Thus, while Δ*nfeR* mutation increases *nfeF* expression and reduces virulence, its mechanism appears to be via subtle effects that are not replicated *in vitro*.

## Conclusions

Despite the urgency of the public health threat posed by classical, multidrug-resistant *K. pneumoniae* strains, our understanding of their mechanisms of infection are still limited. *Galleria mellonella* is an increasingly popular alternative model for bacterial infections, which, unlike mice, is susceptible to infection by c*Kp*. Here, we have performed the first high-throughput fitness profiling study of *K. pneumoniae* during *G. mellonella* infection.

*G. mellonella* had favourable infection parameters for high-throughput screening; the diversity of the highly saturated transposon mutant library was largely maintained through the experiment, and excellent reproducibility was achieved between biological replicates. Infection-related genes identified showed high concordance with current knowledge of *K. pneumoniae* pathogenesis; all of the major virulence factors (siderophores, capsule, O antigen) showed decreased mutant abundance, and many new virulence gene candidates were identified in c*Kp*. This included the metal-dependent regulator *nfeR*, which did not contribute to virulence in the hv*Kp* background. A limitation of our study, and indeed the majority of studies providing molecular detail on *G. mellonella-pathogen* interactions, is that these putative virulence factors have not been further examined in mammalian models.

Our results showed a substantially different fitness landscape for RH201207 and ATCC43816 during *G. mellonella* infection. Fewer infection-related genes were identified in *K. pneumoniae* ATCC43816 (hv*Kp*). This finding may reflect masking of some relevant activities by dominant virulence factors; for example, the highly expressed K2 capsule of this strain may compensate for loss of other cell envelope components, such as multiple genes involved in RpoE signalling which were required in the c*Kp* background. The complex interplay of virulence factors underscores the need to consider the phylogenetic diversity of *K. pneumoniae* when studying its pathogenesis.

We have demonstrated a simple, scalable method for virulence factor profiling in c*Kp*, and used it to provide the first genome-scale view of a c*Kp* infection and compare it to an hv*Kp* strain. The capacity of the *G. mellonella* model to elucidate relevant virulence activities, and the ease with which it can be applied to new strains, opens the possibility for robust species-wide comparisons of infection determinants in *K. pneumoniae* and other opportunistic pathogens.

## Acknowledgements

We thank Matt Mayho, Kim Judge, and the sequencing teams at the Wellcome Sanger Institute for Nanopore and TraDIS sequencing, and the Pathogen Informatics team for support with bioinformatic analysis. We thank Luca Guardabassi and Bimal Jana for providing the RH201207 library.

