## Supplemental Material for "Identifying virulence determinants of multidrug-resistant *Klebsiella pneumoniae* in *Galleria mellonella*"

### Supplementary tables

**Table S1: Bacterial strains, TraDIS libraries, plasmids and primers used in this study**

| **Bacterial strains** | | |
|---|---|---|
| **Strain** | **Description/genotype (Genome accession no)** | **Source** |
| *K. pneumoniae* ATCC 43816 | Hypervirulent, ST493, K-type 2, commonly used in mouse studies (CP009208.1) | Isolate: American Type Culture Collection; Genome: (Broberg *et al.* 2014) |
| *K. pneumoniae* RH201207 | Colistin-resistant UK gut isolate, ST258, K-type 106 | Isolate: (Jana *et al.* 2017); Genome: This study |
| *Escherichia coli* b2163 | $F^-$ RP4-2-Tc::Mu D*dapA*::*(erm-pir)* | (Demarre *et al.* 2005) |
| *K. pneumoniae* RH201207 | Tn5-*rfaH* | (Short *et al.* 2020) |
| *K. pneumoniae* RH201207 | Tn5-*wza* | (Short *et al.* 2020) |
| *K. pneumoniae* RH201207 | Tn5-*wzxE* | This study |
| *K. pneumoniae* RH201207 | Δ*nfeR* (*KPNRH*_00645) | This study |
| *K. pneumoniae* RH201207 | Δ*nfeR* (*KPNRH*_00645) complemented on chromosome | This study |
| *K. pneumoniae* RH201207 | Δ*phoQ* (*KPNRH*_03335) | This study |
| **Transposon mutant libraries** | | |
| **Parent strain** | **Transposon (primers)** | **Source** |
| *K. pneumoniae* RH201207 | Tn5, TetR (Tn5tetR-5Seq, Tn5tetR-5PCR) | (Jana *et al.* 2017) |
| *K. pneumoniae* ATCC 43816 | Tn5-derived from pDS1028 vector, CmR (FS107, FS108) | (Dorman *et al.* 2018) |
| **Plasmids** | | |
| **Name** | **Description** | **Source** |
| pKNG101-Tc | Allelic exchange vector, $Tc^R$ | (Poulter *et al.* 2011) |
| pFLS27 | RH201207 *nfeR* knockout vector, pKNG101-Tc-derived, constructed with FS273-276 | This study |
| pFLS28 | RH201207 *phoQ* knockout vector, pKNG101-Tc-derived, constructed with FS277-280 | This study |
| pFLS36 | RH201207 *nfeR* complementation vector, pKNG101-Tc-derived, constructed with FS273-274 | This study |
| **Primers** | | |
| **Name** | **Sequence 5' – 3'** | **Description** |
| FS107 | GAGCTCGAATTCATCGATGATGGTTGAGATGTGTA | pDS1028 TraDIS 5' Seq |
| FS108 | AATGATACGGCGACCACCGAGATCTACACCAGGAACACTTAACGGCTGACATGG | pDS1028 TraDIS 5' PCR |
| Tn5tetR-5PCR | AATGATACGGCGACCACCGAGATCTACACACTGTGATAAACTACCGCATTAAAGCTTATCG | Tn5-Tet TraDIS 5' PCR |
| Tn5tetR-5Seq | CGATGATAAGCTGTCAAACATTGATGGTTGAGATGTGTA | Tn5-Tet TraDIS 5' Seq |
| FS273 | gctACTAGTTGCCCTACATTGAAGATGC | Mutant construction, SpeI site |

| | | |
|---|---|---|
| FS274 | gctACTAGTCGTAAGCCAGCTCGTGA | Mutant construction, SpeI site |
| FS275 | GGTGGTGAAAAAAGAGGCGGTTACCTCCTGCTGT TTT | Mutant construction, overlap |
| FS276 | ACAGCAGGAGGTAACCGCCTCTTTTTTCACCACC GGC | Mutant construction, overlap |
| FS277 | gctACTAGTCGGGCAGTGCTGTTTCA | Mutant construction, SpeI site |
| FS278 | gctACTAGTGCGAACATCTCCCGGAT | Mutant construction, SpeI site |
| FS279 | GTGAAAAATCTCAACGACAGCGCAATTCGAACA GAT | Mutant construction, overlap |
| FS280 | TCGAATTGCGCTGTCGTTGAGATTTTTCACGGCG | Mutant construction, overlap |

**Table S2: TraDIS sequencing information**

| Strain | Sample | Sample Acc | Lane 1 | Lane 2 | Run 1 | Run 2 | Sample Read Count | Transposon Reads | Mapped Reads | Unique Insertions |
|---|---|---|---|---|---|---|---|---|---|---|
| RH201207 | *G. mellonella* infection Input 1 | ERS2856142 | 24686_1#10 | 24686_2#10 | ERR3313543 | ERR3313564 | 14,671,625 | 14,611,926 | 13,067,937 | 426,116 |
| RH201207 | *G. mellonella* infection Input 2 | ERS2856143 | 24686_1#11 | 24686_2#11 | ERR3313544 | ERR3313565 | 15,139,931 | 15,075,494 | 13,489,139 | 432,678 |
| RH201207 | *G. mellonella* infection Input 3 | ERS2856144 | 24686_1#12 | 24686_2#12 | ERR3313545 | ERR3313566 | 13,907,983 | 13,848,936 | 12,399,651 | 426,248 |
| RH201207 | *G. mellonella* infection 2 hpi 1 | ERS2856148 | 24686_1#16 | 24686_2#16 | ERR3313549 | ERR3313570 | 14,008,792 | 13,952,271 | 12,471,099 | 254,076 |
| RH201207 | *G. mellonella* infection 2 hpi 2 | ERS2856149 | 24686_1#17 | 24686_2#17 | ERR3313550 | ERR3313571 | 14,494,416 | 14,430,166 | 12,887,292 | 256,592 |
| RH201207 | *G. mellonella* infection 2 hpi 3 | ERS2856150 | 24686_1#18 | 24686_2#18 | ERR3313551 | ERR3313572 | 14,543,107 | 14,481,084 | 12,959,253 | 256,600 |
| RH201207 | *G. mellonella* infection 6 hpi 1 | ERS2856151 | 24686_1#19 | 24686_2#19 | ERR3313552 | ERR3313573 | 14,752,537 | 14,691,081 | 13,174,812 | 254,756 |
| RH201207 | *G. mellonella* infection 6 hpi 2 | ERS2856152 | 24686_1#20 | 24686_2#20 | ERR3313553 | ERR3313574 | 14,053,700 | 13,996,625 | 12,594,311 | 211,200 |
| RH201207 | *G. mellonella* infection 6 hpi 3 | ERS2856153 | 24686_1#21 | 24686_2#21 | ERR3313554 | ERR3313575 | 13,031,469 | 12,975,166 | 11,619,148 | 233,170 |
| ATCC 43816 | *G. mellonella* infection Input 1 | ERS2856117 | 23257_1#1 | 23363_1#1 | ERR4706232 | ERR4706238 | 1,966,025 | 1,959,181 | 1,901,817 | 301,651 |
| ATCC 43816 | *G. mellonella* infection Input 2 | ERS2856118 | 23257_1#2 | 23363_1#2 | ERR4706233 | ERR4706239 | 1,858,337 | 1,851,902 | 1,796,586 | 301,089 |
| ATCC 43816 | *G. mellonella* infection Input 3 | ERS2856119 | 23257_1#3 | 23363_1#3 | ERR4706234 | ERR4706240 | 2,003,569 | 1,996,407 | 1,937,781 | 307,248 |
| ATCC 43816 | *G. mellonella* infection 4 hpi 1 | ERS2856120 | 23257_1#4 | 23363_1#4 | ERR4706235 | ERR4706241 | 2,085,144 | 2,076,503 | 2,001,084 | 234,563 |
| ATCC 43816 | *G. mellonella* infection 4 hpi 2 | ERS2856121 | 23257_1#5 | 23363_1#5 | ERR4706236 | ERR4706242 | 1,962,113 | 1,953,683 | 1,882,388 | 227,021 |
| ATCC 43816 | *G. mellonella* infection 4 hpi 3 | ERS2856122 | 23257_1#6 | 23363_1#6 | ERR4706237 | ERR4706243 | 1,945,318 | 1,938,289 | 1,875,950 | 232,356 |

**Table S3: Pearson correlation coefficient $r^2$ values of the comparisons of biological replicates by reads per gene, insertion indices per gene and reads per unique insertion site (UIS), respectively.**

| Sample | Reads per gene | | | Insertion index per gene | | | Reads per UIS | | |
|---|---|---|---|---|---|---|---|---|---|
| | rep 1 vs rep 2 | rep 2 vs rep 3 | rep1 vs rep 3 | rep 1 vs rep 2 | rep 2 vs rep 3 | rep1 vs rep 3 | rep 1 vs rep 2 | rep 2 vs rep 3 | rep1 vs rep 3 |
| **RH201207 input** | 0.9988 | 0.9988 | 0.9976 | 0.9943 | 0.9945 | 0.9944 | 0.9589 | 0.959 | 0.9548 |
| **RH201207 2 hpi** | 0.948 | 0.9957 | 0.995 | 0.983 | 0.984 | 0.9831 | 0.7692 | 0.769 | 0.7704 |
| **RH201207 6 hpi** | 0.9922 | 0.9938 | 0.9938 | 0.9801 | 0.9792 | 0.9828 | 0.7078 | 0.6929 | 0.7337 |
| **ATCC 43816 input** | 0.9983 | 0.9982 | 0.9979 | 0.9845 | 0.9838 | 0.9852 | 0.9627 | 0.9644 | 0.9628 |
| **ATCC 43816 4 hpi** | 0.9959 | 0.9941 | 0.9924 | 0.9764 | 0.9765 | 0.9772 | 0.9145 | 0.913 | 0.9151 |

### Supplementary figures

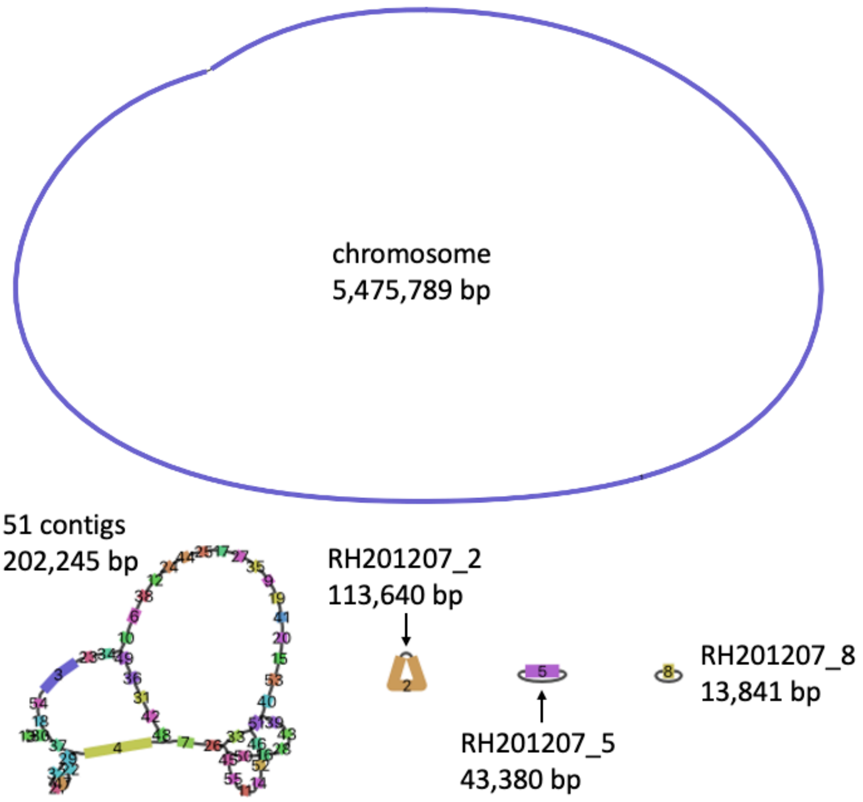


**Figure S1: Assembly graph of the RH201207 genome.** The genome was assembled using long Nanopore and short Illumina reads using Unicycler. This graphical representation of the assembly was generated with Bandage and shows a circularised chromosome and three circularised plasmids. One structure, with a length of 202,245 bp, probably representing one or multiple plasmids could not be fully resolved.

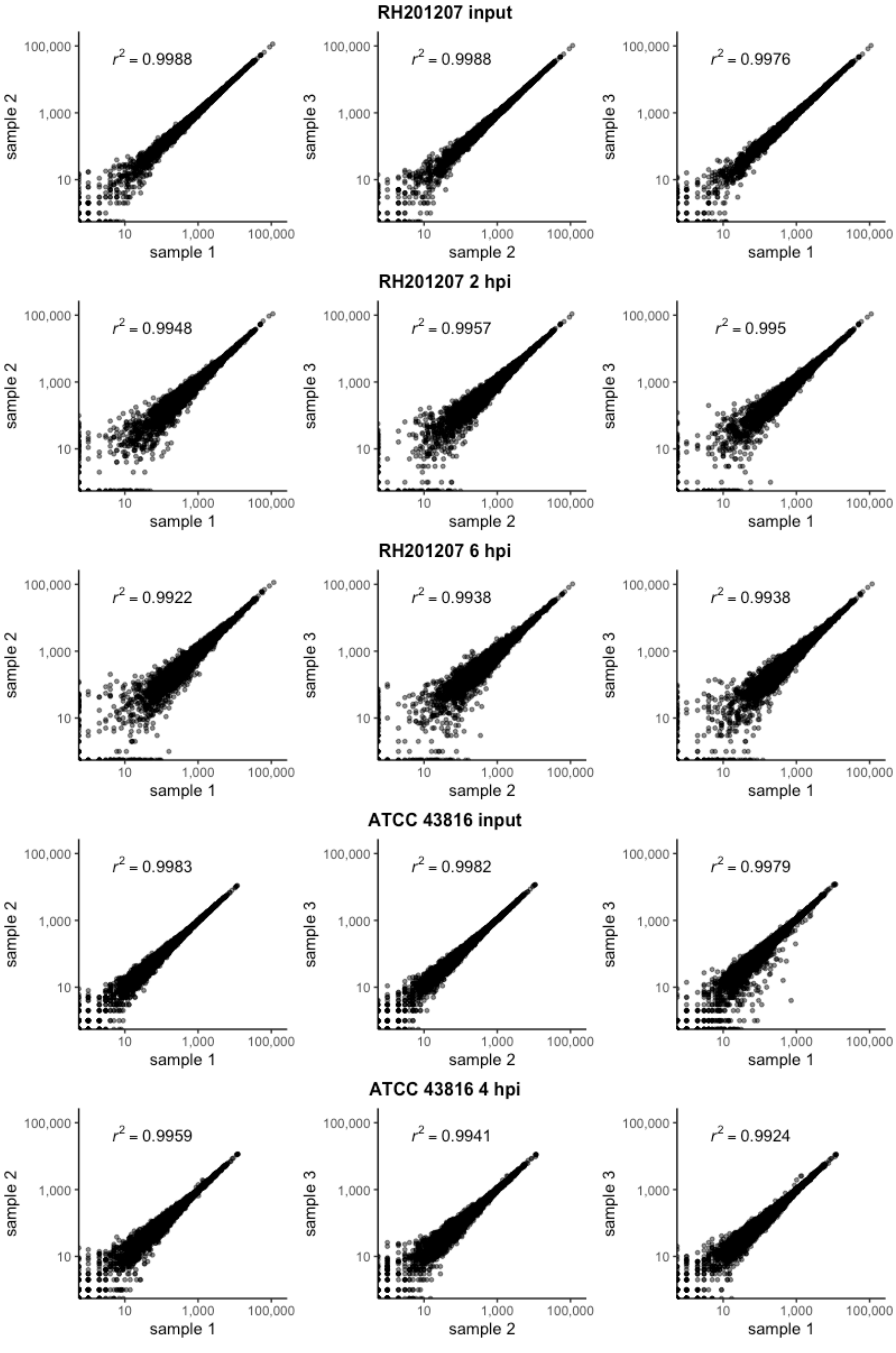


**Figure S2: Pairwise correlation of reads per gene of biological TraDIS replicates and their Pearson correlation coefficients**

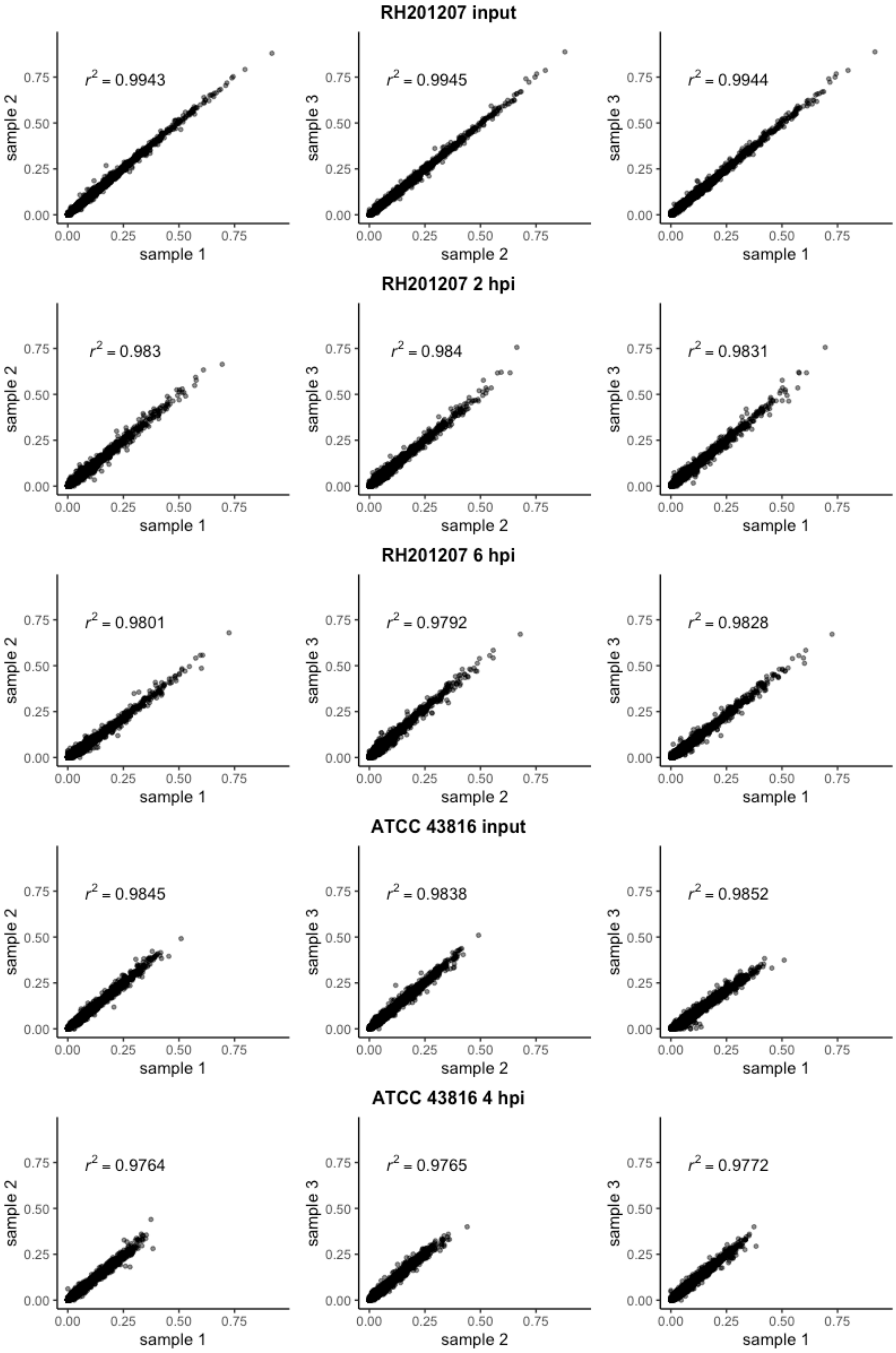


**Figure S3: Pairwise correlation of insertion indices per gene of biological TraDIS replicates and their Pearson correlation coefficients**

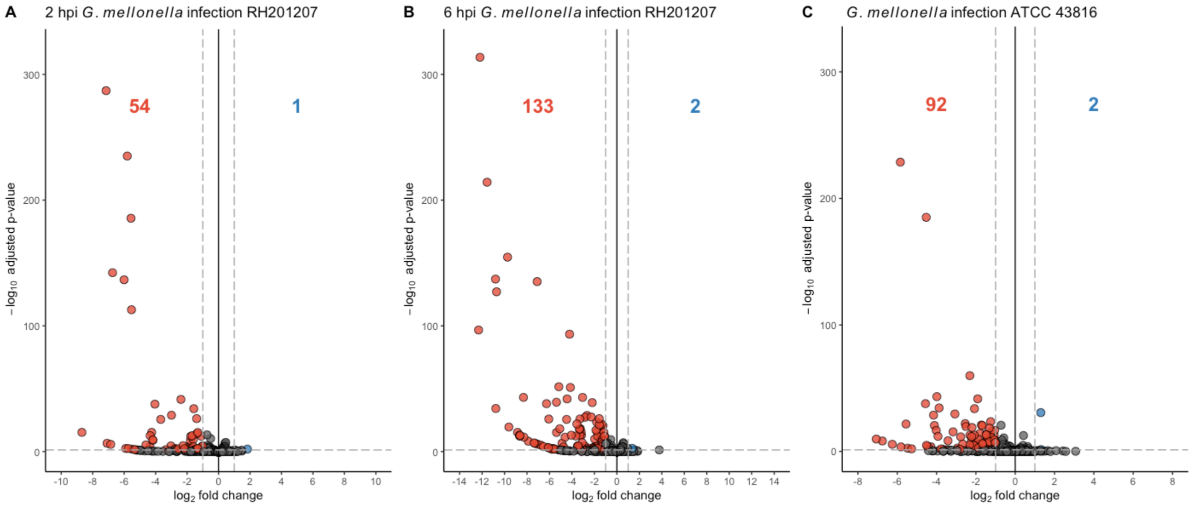


**Figure S4: Volcano plots of TraDIS datasets.** Significantly less abundant genes after infecting *G. mellonella* larvae are shown in red and significantly higher abundant genes in blue. Non-significant genes are shown in grey. Genes were considered significant if they possessed a Benjamini-Hochberg corrected *p*-value below a threshold of 0.05 and an absolute log2 fold-change greater than 1. Essential and ambiguous-essential genes are removed from the analysis. A: RH201207 at 2 hpi, B: RH201207 at 6 hpi, C: ATCC 43816 at 4 hpi.

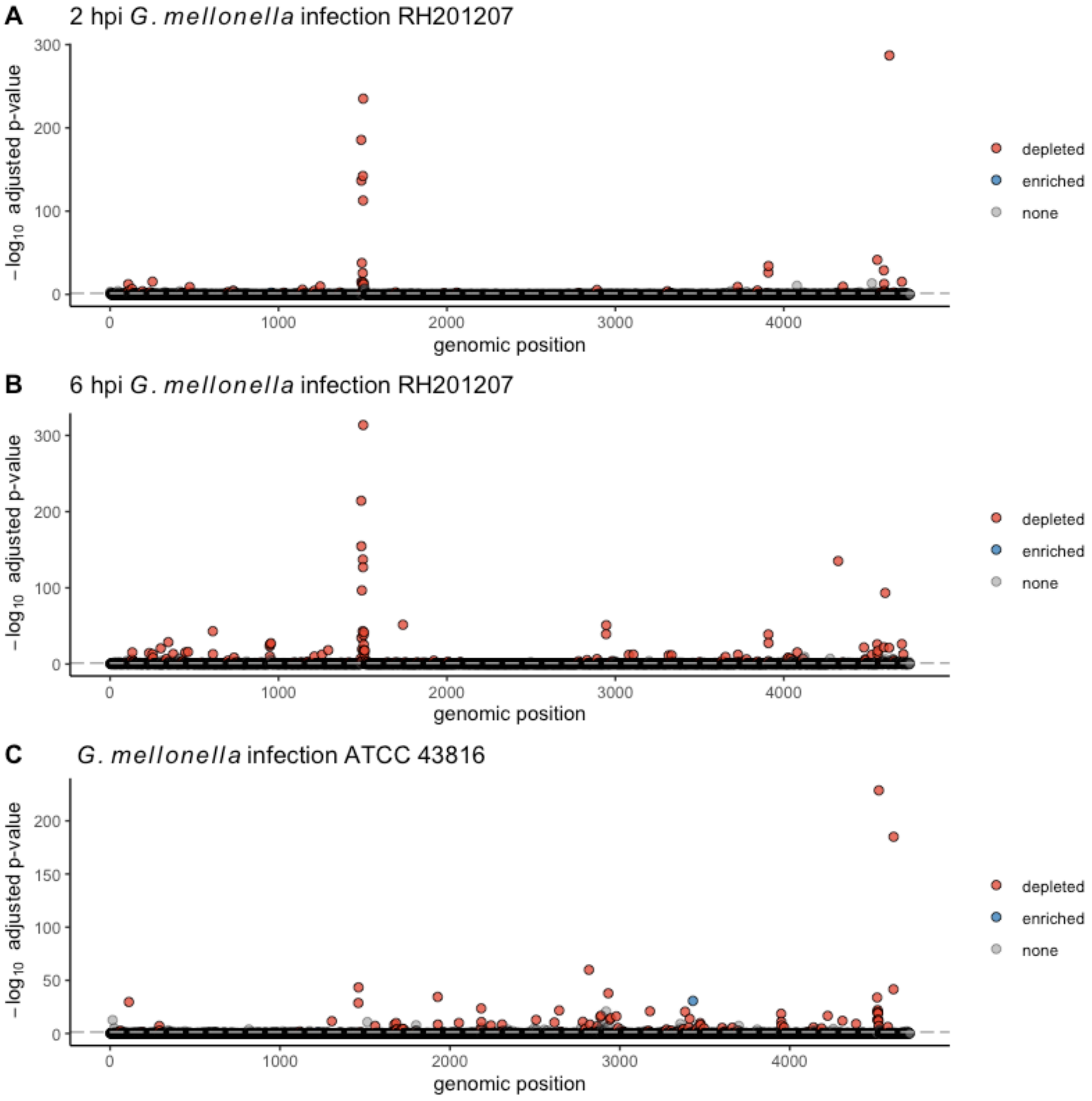


**Figure S5: Manhattan plots of TraDIS datasets showing adjusted p-value vs genomic position.** Significantly less abundant genes after infecting *G. mellonella* larvae are shown in red and significantly higher abundant genes in blue. Non-significant genes are shown in grey. Genes were considered significant if they possessed a Benjamini-Hochberg corrected *p*-value below a threshold of 0.05 and an absolute log2 fold-change greater than 1. Essential and ambiguous-essential genes are removed from the analysis. A: RH201207 at 2 hpi, B: RH201207 at 6 hpi, C: ATCC 43816 at 4 hpi.

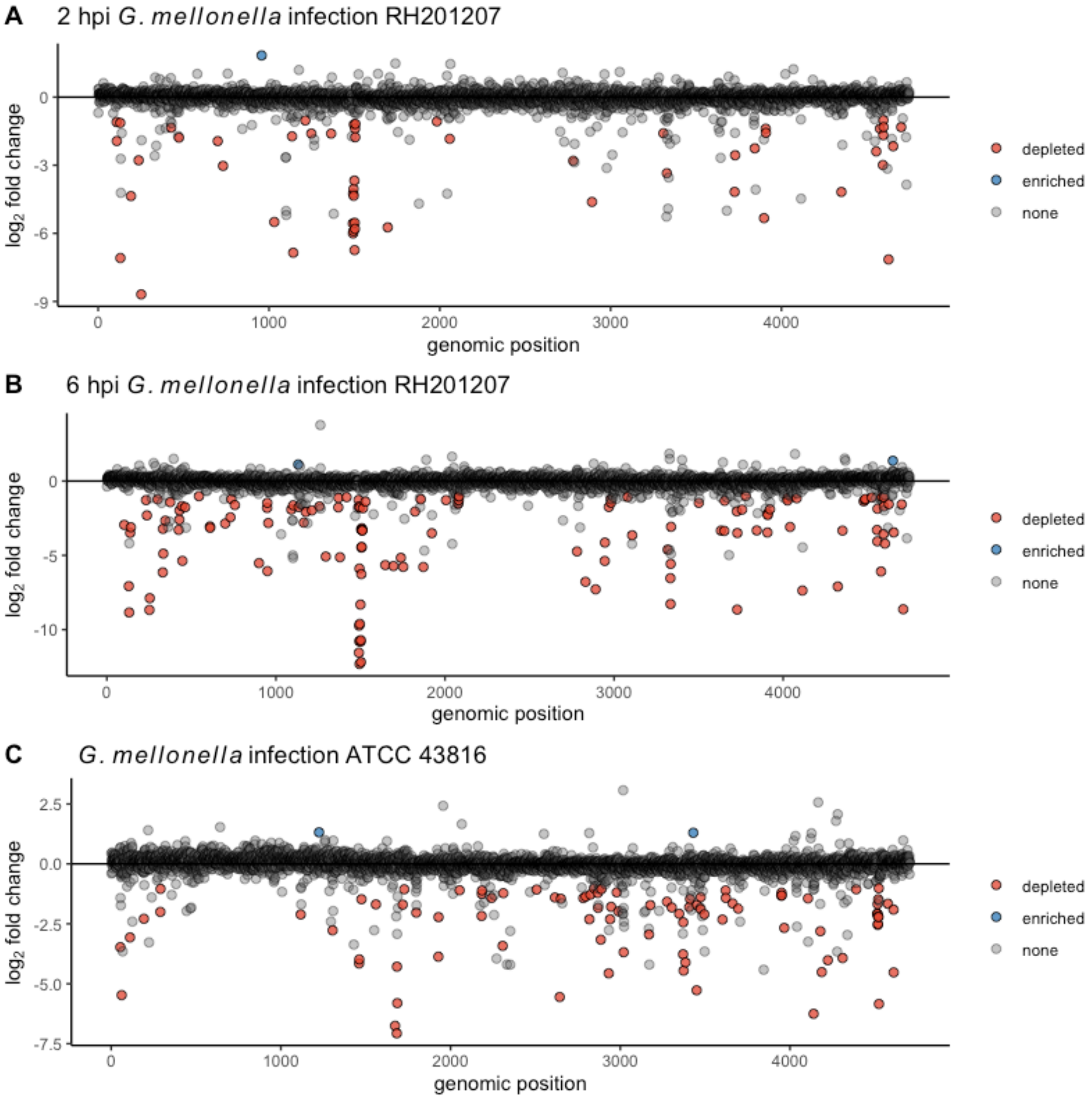


**Figure S6: Manhattan plots of TraDIS datasets showing log2 fold-change vs genomic position.** Significantly less abundant genes after infecting *G. mellonella* larvae are shown in red and significantly higher abundant genes in blue. Non-significant genes are shown in grey. Genes were considered significant if they possessed a Benjamini-Hochberg corrected *p*-value below a threshold of 0.05 and an absolute log2 fold-change greater than 1. Essential and ambiguous-essential genes are removed from the analysis. A: RH201207 at 2 hpi, B: RH201207 at 6 hpi, C: ATCC 43816 at 4 hpi.
